# A microbial perspective on balancing trade-offs in ecosystem functions in a constructed stormwater wetland

**DOI:** 10.1101/2020.04.01.020776

**Authors:** Regina B. Bledsoe, Eban Z. Bean, Samuel S. Austin, Ariane L. Peralta

**Affiliations:** East Carolina University, Department of Biology, Greenville, NC, 27858, USA; University of Florida, Department of Agricultural and Biological Engineering, Gainesville, FL, 32611, USA

**Keywords:** Greenhouse gases, Denitrification, Methane, Stormwater, Urban ecosystems

## Abstract

Green stormwater infrastructure, such as constructed wetlands (CWs), is a type of stormwater control measure that can decrease nutrient and pollutant loads from urban stormwater runoff. Wetland soil microorganisms provide nutrient and pollutant removal benefits which can also result in ecosystem disservices such as greenhouse gas (GHG) emissions and can inadvertently exacerbate climate change. Microbial respiration by facultative anaerobes in anoxic conditions is the primary pathway for nitrogen removal (benefit). Similar anoxic conditions that support denitrifying microorganisms can also support obligate anaerobes that produce methane (CH_4_) via methanogenesis (disservice). We examined nitrogen removal potential, GHG production, and microbial community structure within permanently flooded and shallow land or temporarily-flooded areas of a stormwater CW to identify zones for CW design optimization. Results indicate that permanently flooded zones compared to shallow land zones are greater sources of CH_4_ emissions (80.80 ± 118.31, 2.32 ± 9.33 mg CH_4_-C m^-2^ hr^-1^, respectively) and emit more carbon to the atmosphere (7161.27 kg CO_2_, 93.20 kg CO_2_ equivalents, respectively). However, nitrogen removal potential rates were similar across both flooded and shallow land zones (24.45 ± 20.18, 20.29 ± 15.14 ng N_2_O-N hr^-1^ g^-1^ dry soil, respectively). At this particular CW, reduction of permanently flooded zones within the wetland could decrease GHG emissions (disservice) without limiting nitrogen removal (benefit) potential of the wetland. Holistic development and design of stormwater control measures, which account for microbial activity, provides the opportunity to maximize benefits (i.e., nutrient and pollutant removal) and reduce disservices (i.e., GHG emissions) of green stormwater infrastructure.

## Introduction

In urban areas, impervious surfaces increase nutrient and pollutant loads in stormwater runoff, thereby decreasing water quality within neighboring watersheds (Bell et al., 2019; Line and White, 2007; O’Driscoll et al., 2010). Nutrient loadings are of particular concern due to adverse effects (e.g., harmful algal blooms and hypoxic zones) on downstream water quality (Paerl et al., 2014; Rabalais et al., 2009). Green stormwater infrastructure, such as constructed wetlands (CWs), are stormwater control measures (SCMs) that are designed to mimic natural ecosystems to provide relief from stormwater runoff and improve water quality in and around urban areas. The physical, chemical, and biological treatment processes that naturally occur in wetlands make CWs an effective approach to reduce nutrient loads in stormwater runoff (Koch et al., 2014; Mitsch et al., 2013; Payne et al., 2014; Reisinger et al., 2016). Thus, entities such as the North Carolina Department of Environmental Quality encourage use of SCMs as a way to reduce nutrient loadings to waterways by offering nutrient reduction credits. CWs that have a ponding depth of less than 12 inches and a hydraulic residence time of two to five days are credited with 40% removal of nitrogen (N) from stormwater runoff (NCDEQ, 2018). However, this nutrient reduction policy and similar policies outside of North Carolina do not consider the potential trade-off between water quality (e.g., N removal) and air quality (e.g., greenhouse gas (GHG) emissions). The unintended GHG production from stormwater CWs could negatively affect climate change mitigation and adaptation by reducing the carbon storage capacity typically provided by wetlands (Demuzere et al., 2014). Therefore, in order to meet management goals in a more holistic way (i.e., enhancing total N removal and reducing GHG emissions), a deeper understanding of microbial processes occurring within CWs are needed to optimize wetland design.

Plants and soil microorganisms carry out nutrient and pollutant removal processes that occur within wetlands. While plants and microorganisms can assimilate ammonium (NH_4_^+^) and nitrate (NO_3_^-^) ions into biomass, the primary pathway of N removal from a wetland ecosystem is microbial respiration via denitrification and to a lesser degree anaerobic ammonium oxidation (Lee et al., 2009). Denitrification is typically coupled with nitrification in which NH_4_^+^ is first transformed to nitrite (NO_2_^-^) and then NO_3_^-^ by aerobic nitrifying microorganisms (Lee et al., 2009). The process of denitrification occurs when NO_3_^-^ is reduced to nitric oxide (NO), then to nitrous oxide (N_2_O), a potent GHG, and finally to dinitrogen (N_2_), an inert and abundant atmospheric gas (Knowles, 1982; Lee et al., 2009; Smith and Tiedje, 1979). Denitrification is an anaerobic microbial process that requires anoxic or low-oxygen conditions such as those found in saturated and flooded soils within wetlands (Marton et al., 2015; Smith and Tiedje, 1979). Many denitrifying microorganisms are facultatively anaerobic, meaning these microorganisms can perform aerobic respiration when oxygen is available but can use other alternate electron acceptors (i.e., nitrate) when anoxic soil conditions arise (Tiedje, 1988). Due to this facultative anaerobic metabolism, these microorganisms respond to hydrologic change with varying degrees depending on past environmental conditions (Peralta et al., 2014, 2013).

Similar anoxic conditions that support nitrogen removal by denitrification (transformation of NO_3_^-^ to N_2_) can also support production of methane (CH_4_), a GHG, which is considered an ecosystem disservice (Demuzere et al., 2014). Methanogens are obligate anaerobes that produce CH_4_ via methanogenesis but only under anoxic conditions when the environment is depleted of other electron acceptors (e.g., nitrate, sulfate, and ferric iron) (Fetzer and Conrad, 1993; Liu et al., 2008). When organic matter is decomposed by fermentation, methanogens can use the byproducts of acetate, hydrogen, and CO_2_ for energy and produce CH_4_ (Conrad, 2007). CH_4_ production can also support the consumption of CH_4_ by aerobic methanotrophs, which have been reported to consume 45-90% of CH_4_ produced (Brindha and Vasudevan, 2018; Conrad, 2007; Le Mer and Roger, 2001). Systems in which the population of methanogens (i.e., producers of CH_4_) is greater than methanotrophs (i.e., consumers of CH_4_) can lead an ecosystem to be a CH_4_ source (Altshuler et al., 2019; Rey-Sanchez et al., 2019; Wen et al., 2018). Since methanogens are obligate anaerobes, even low amounts of oxygen can suppress CH_4_ production (Fetzer and Conrad, 1993; Le Mer and Roger, 2001). However, methanotrophs are aerobes, and there is typically greater CH_4_ oxidation in dynamic, wet-dry conditions that are not permanently inundated (Chowdhury and Dick, 2013). Therefore, anoxic conditions that can support denitrification can also result in GHG emissions, especially when nitrate is limiting (Burgin et al., 2011).

The three major biogenic GHGs of concern, CO_2_, CH_4_, and N_2_O, are known to contribute to increased atmospheric warming and thereby influence global climate change. While CO_2_ (416.21 ppm) is the most abundant GHG in the atmosphere, CH_4_ (1.87 ppm) and N_2_O (0.33 ppm) are found in lower atmospheric concentrations but have a greater global warming potential (GWP) per molecule than CO_2_ (IPCC, 2014) (https://www.esrl.noaa.gov/gmd/ccgg/trends/). Specifically, CH_4_ and N_2_O have GWP of 28 and 265, respectively, over a 100 year time-scale when compared to CO_2_ which is equal to 1 (IPCC, 2014).Therefore, smaller amounts of CH_4_ or N_2_O can contribute more to increasing GWP than equal amounts of CO_2_.

To improve water quality management of CWs in the urban landscape, both ecosystem services (e.g., nitrogen removal by denitrification) and disservices (e.g., GHG production) should be considered during the planning and design phase (Demuzere et al., 2014). To accommodate stormwater runoff and water quality functions, CWs generally have four main zones: upland (rarely submerged except during large runoff events), shallow land (soil is saturated but only temporarily inundated following runoff events), shallow pools (designed to be permanently inundated except during droughts) and deep pools (permanently inundated) (NCDEQ, 2019). In CWs, ammonification and nitrification occur in the shallow land zones as stormwater percolates into aerated soils. If soils in shallow lands are saturated for sufficient durations, denitrification is expected to eventually occur as well. It is in the shallow water zones where denitrification is conventionally thought to occur due to saturated, anoxic conditions. In this study we focus on denitrification as the ecosystem service since this process typically accounts for the most N removal in CWs (Rahman et al., 2019).

The goal of this study is to identify areas within a stormwater CW that support denitrification and reduce GHG emissions, especially due to methanogenesis. We hypothesize that the dynamic wet-dry conditions occurring in shallow land and flooded zones will both support the anaerobic process of denitrification, while only the flooded zones will be sources of CH_4_ gas. If production of CH_4_ is greater than CO_2_ production, the overall GWP of the wetland will increase since CH_4_ has a higher GWP than CO_2_. However, due to the facultative nature of microbes residing in the shallow land zones, denitrification will still occur in shallow land zones after a short period of flooding if NO_3_^-^ is not limiting. Further, we expect microbial community composition to differ between land types such that flooded zones will be represented by a greater relative abundance of microbes involved in CH_4_ cycling while shallow land zones will have a greater proportion of microbes involved in nitrogen cycling.

## 2. Materials and methods

To test these hypotheses, we monitored GHG fluxes monthly for one year and sampled sediments and surface water for chemical analyses seasonally. To determine microbial (bacterial and archaeal) community structure in sediments, we carried out 16s rRNA V4 amplicon sequencing. We also determined potential denitrification rates across the wetland using a short-term incubation experiment. By combining GHG rates with potential denitrification rates, sediment and water chemistry parameters along with microbial community composition, we examined the potential benefits and disservices occurring within this CW.

### 2.1 Study site

The study location is a recently constructed (2015) stormwater wetland located on East Carolina University’s main campus in Greenville, NC (N 35°35’22.4”, W 77°22’13.4”). Prior to construction, the area had served as a dry detention basin since at least 1998, based on historical aerial imagery analysis. The 0.13 acre CW receives runoff from 5.7 acre parking lot in a highly urbanized area of the city. To create the additional storage volume for runoff and permanent ponding, the basin surface was excavated to create landform features. Soils were characterized by orange and gray mottled loamy clay, indicative of reducing, low-oxygen soil conditions. Infiltration rates were measured to be < 0.1 in/h, making for a suitable substrate to maintain the permanent water levels in the CW. The temporary ponding depth (30.5 cm) of the wetland is equal to 0.51 cm of runoff within the impervious drainage area (Fig. 1), undersized relative to permitting requirements at the time (NCDEQ, 2019). The total surface area is divided among main zones (% surface area): shallow land (35%), shallow water (35%), deep pools (10%), forebay (10%), and outlet pool (10%) (NCDEQ, 2019).The dominant plants within saturated areas of the wetland are *Typha* spp., *Pontederia cordata*, and *Iris versicolor* L. The shallow land area along the water’s edge is dominated by *Carex lupulina* Muhl. ex Willd. 2.2 *GHG monitoring and flux calculations*

**Figure 1.**
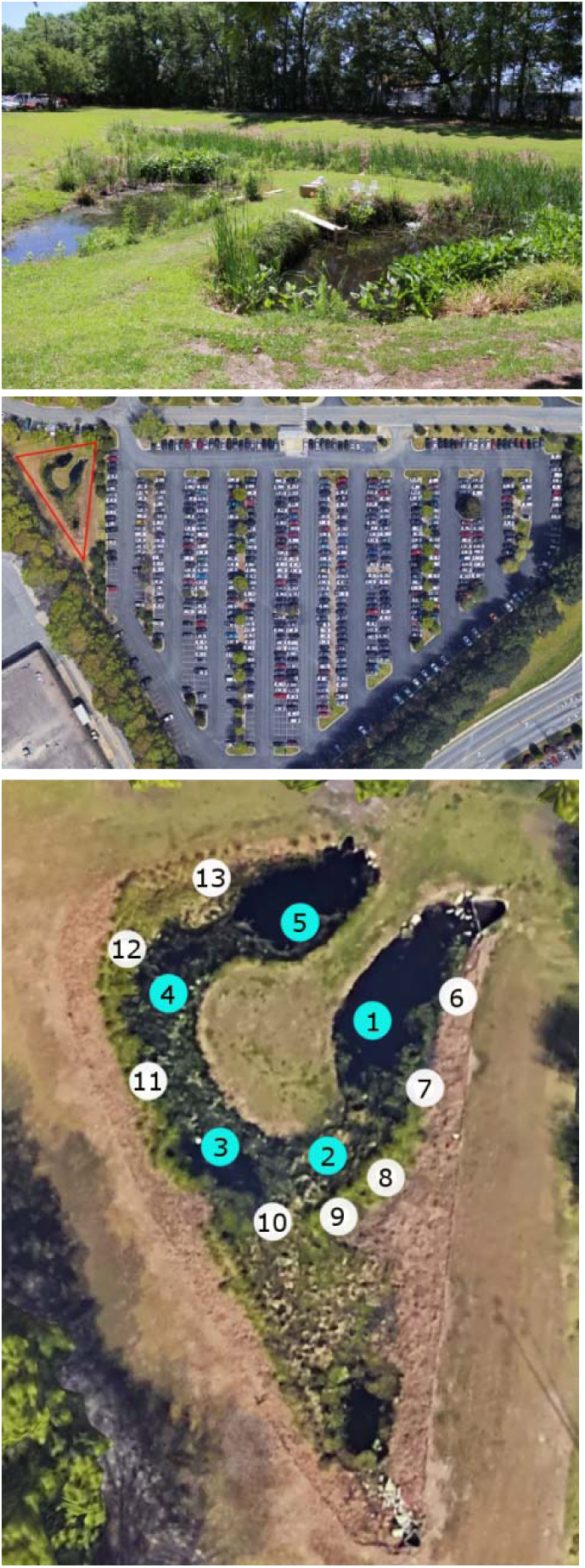
Top: Landscape photograph of wetland. Center: Google Earth image of parking lot and stormwater CW outlined in red. Bottom: Google Earth image of wetland with blue circles (1-5) representing flooded plots and white circles (6-13) representing shallow land plots sampled for GHG and sediments.

We used a static chamber method to measure soil respiration as GHG fluxes (Parkin et al., 2010). At the CW, we established 13 GHG sampling plots along two main transects that follow the central channel within the wetland that flows from inlet to outlet (Fig. 1). Of those 13 plots, chambers 1-5 were placed in shallow water and deep pools (flooded zones), and chambers (6-13) were placed in shallow land zones along the outside edge of the main channel. Each sampling plot in shallow land zones (n = 10) was fitted with a GHG chamber bottom installed 6 cm belowground. During sampling events, chamber tops (30 cm height, 15 cm diameter) were attached to an aboveground groove, and the groove was filled with water to create a gas tight seal. GHG chamber tops in flooded zones were fitted with a Styrofoam float allowing 2-3 cm of the chamber top below the water surface.

We collected GHG samples monthly for one year. Sampling events occurred at least 48 hours after a rain event of > 0.51 cm. Gas samples were collected at 0, 12, 24, and 36 minutes using 20 mL syringes. Each gas sample was divided between two 3.7 mL Exetainers® (Labco, Lampeter, Wales, UK) with dual septa and stored upside at room temperature until analyzed. We measure GHG concentrations using a Shimadzu 2014 Gas Chromatograph (Shimadzu Scientific Instruments, Columbia, Maryland) fitted with an electron capture detector to detect N_2_O and flame ionization detector to measure CH_4_ and CO_2_. Monthly GHG fluxes were calculated using the linear change in concentration and the ideal gas law (Millar et al. 2018). Seasonal estimates were calculated by averaging monthly flux measurements: spring = March-May, summer = June-August, fall = September-November and winter = December-February. In addition, the equivalent CO_2_ flux for CH_4_ was calculated by multiplying CH_4_ flux by 28, which is equal to the increase in warming potential in one molecule of CH_4_ versus one molecule of CO_2_.

### 2.4 Soil collection and chemical properties

We characterized soil physicochemical properties seasonally (May, July, October, and December). Each composite soil sample represented three cores (3.1 cm diameter, 12 cm depth) that were adjacent to each GHG sampling chamber. We homogenized each composite sample by passing the sample through a 4 mm sieve, and then processed subsamples for soil gravimetric moisture, pH, extractable ammonium and nitrate, total carbon, and total nitrogen. We combined 5 g of field moist soil and 45 mL of 2M potassium chloride and shook samples for 1 hour before gravity filtration to collect soil extracts. NH_4_^+^ and NO_3_^-^ concentrations were colorimetrically measured using a SmartChem 200 auto analyzer (Unity Scientific Milford, Massachusetts, USA) at the East Carolina University Environmental Research Laboratory. Using a 20-30 g subsample, we measured field-moist and oven-dry (dried 24 hours at 105°C) soil weights to determine moisture content by dividing the mass of water by the mass of oven-dried soil. Using a mortar and pestle, we coarsely ground a subsample of soil and combined 10 g of soil with 10 mL of Nanopure® water to measure soil pH. On another subsample, we finely ground the soil for elemental carbon and nitrogen analysis using an elemental analyzer (2400 CHNS Analyzer; Perkin Elmer; Waltham, Massachusetts, USA) at the Environmental and Agricultural Testing Service laboratory (Department of Crop and Soil Sciences at NC State). Finally, we measured organic matter content based on the loss on ignition method by ashing 20 g coarsely ground oven-dried soil at 550 °C (Hoogsteen et al., 2015) for 2 hours since the majority of OM is ashed within the first 2 hours of combustion (Heiri et al., 2001). However, up to 4 hours may be required to completely ash all organic matter from soils with high OM content (Heiri et al., 2001). In addition, a subsample of field moist soil was stored at 4 °C until we performed denitrification potential assays. Finally, a subsample of soil was stored at -80 °C until DNA extraction for bacterial amplicon sequencing.

### 2.3 Seasonal denitrification potential

Within 72 hours of soil collection, we measured potential denitrification rates using the acetylene block denitrification enzyme assay method (Schaller et al., 2004; Tiedje et al., 1989; Wall et al., 2005). We weighed triplicate soil subsamples of 25 g and transferred each subsample into separate 125 mL Wheaton bottles fitted with a phenolic cap and butyl septa. In each bottle, we added 75 mL of 1 mM potassium chloride (KNO_3_) to provide an abundant nitrogen supply and 1.3 mL of chloramphenicol (100 mg/mL) to inhibit new denitrification enzyme production. Bottles were sealed with septa-centered caps, shook, and purged with helium for 5 minutes to create an anoxic environment. We removed 15 mL of helium from the headspace and replaced with 15 mL of pure acetylene gas to prevent the conversion of N_2_O to N_2_. At each hour for 3 hours, we collected a 10 mL gas sample (time point T0, T1, T2, T3) from each bottle and stored the gas sample in a 3.7 mL Exetainer®. We shook each bottle to equilibrate N_2_O in aqueous and sediment phases before each sample collection. We added 10 mL of 10% acetylene mixture to the headspace after each sample collection to maintain acetylene concentrations within the bottle. N_2_O fluxes were calculated as described for GHG fluxes (Millar et al., 2018).

### 2.5 Microbial Community Analysis

We examined the microbial community from soils collected in May (spring) and October (fall) from flooded plots (1-5) and shallow land plots (7-9, 11-13) (Fig. 1). First, we extracted genomic DNA from soils using the Qiagen DNeasy Powerlyzer PowerSoil Kit. Genomic DNA was amplified using barcoded primers 515FB/806R primer set, originally developed by the Earth Microbiome Project (earthmicrobiome.org) to target the V4-V5 region of the bacterial 16S subunit of the ribosomal RNA gene (Apprill et al., 2015; Caporaso et al., 2010; Parada et al., 2016). Samples were prepared for sequencing by using polymerase chain reaction (PCR) chemistry to amplify the 16S rRNA DNA with a unique sample identifier to each sequence within each individual sample. For each sample, three 50 µL PCR libraries were prepared by combining 38.35 µL molecular grade water, 5 µL Amplitaq Gold 360 10x buffer, 2.4 µL MgCl_2_ (25 mM), 1 µL dNTPs (40mM total, 10mM each), 0.25 µL Amplitaq Gold 360 polymerase, 1 µL 515 forward barcoded primer (10 µM), 1 µL 806 reverse primer (10 µM), and 1 µL DNA template (10 ng µL^-1^) following the manufacturer’s protocol (Applied Biosystems, Foster City, CA, USA. Thermocycler conditions for PCR reactions were as follows: initial denaturation (94 °C, 3 minutes); 30 cycles of 94°C for 45 seconds, 50 °C for 30 seconds, 72 °C for 90 seconds; final elongation (72 °C, 10 minutes). The triplicate 50 µL PCR libraries were combined and then amplified DNA sequences separated from PCR reagents using the AMPure XP magnetic bead protocol (Axygen, Union City, California, USA). Cleaned PCR products were quantified using QuantIT dsDNA broad range assay (Thermo Scientific, Waltham, Massachusetts, USA) and diluted to a concentration of 10 ng µL^-1^. We combined barcoded PCR samples (5 ng µL^-1^) in equimolar concentration and sequenced the pooled library using the Illumina MiSeq platform paired end read approach (Illumina Reagent Kit v2, 500 reaction kit) at the Indiana University Center for Genomics and Bioinformatics Sequencing Facility.

Sequences were processed using a standard mothur pipeline (v1.40.1)(Kozich et al., 2013; Schloss et al., 2009). We assembled contigs from the paired end reads, quality trimmed using a moving average quality score (minimum quality score 35), aligned sequences to the Silva Database (Quast et al. 2013; SSURef v123), and removed chimeric sequences using the VSEARCH algorithm (Rognes et al., 2016). We created operational taxonomic units (OTUs) by first splitting sequences based on taxonomic class and then binning into OTUs based on 97% sequence similarity. The Silva database was used to assign taxonomy to each OTU.

### 2.6 Statistical Analysis

All statistical analyses were completed in the R statistical environment (RStudio v1.1.383, Rv3.4.0) (R Core Team, 2019). We constructed linear mixed effects models with sampling plot as a random effect using the *lme4* R package (Bates et al., 2015) to determine the importance of hydrology, season, and distance from the inlet, and the interactions on GHG fluxes and denitrification potential fluxes within the constructed wetland. We then conducted model comparisons using sample size corrected Akaike information criteria (AICc) model comparisons, which adjust for small sample size, against the null model to determine which fixed effects (hydrology,season, and distance from inlet) to determine which combinations had the most explanatory power (Gorsky et al., 2019; Hurvich and Tsai, 1993). Estimated marginal means with Tukey post-hoc adjustment in the R package *emmeans* (Lenth, 2017) was used to determine significance (p<0.05) in gas flux rates between treatment groups and the *multcomp* R package (Hothorn et al., 2020) was used to assign Tukey post-hoc groups. Lastly, we used the *MuMIn* R package to determine the proportion of variance explained by fixed effects (marginal) and the complete model (conditional) (Barton, 2019; Gorsky et al., 2019). The CH_4_, CO_2_, and N_2_O fluxes were transformed to the log of the cube root and denitrification potential rates were log transformed to meet normality assumptions of the statistical models.

Finally, we examined microbial community diversity and how that relates to soil chemistry and GHG fluxes. We standardized read counts before calculating diversity metrics for sediment microbial communities by removing singletons and doubletons, rarifying all sample to the sample with the lowest read count, and resampling. Principal Coordinates of Analysis (PCoA) based on Bray-Curtis dissimilarity was used to visualize patterns of bacterial diversity among treatments and sampling dates. Permuted analysis of variance (PERMANOVA) was run using the *adonis* function in the *vegan* package (Oksanen, 2015) to determine the importance of hydrology and season in shaping overall microbial community composition. We performed a Dufrene-Legendre indicator species analysis using the *labdsv* R package (Roberts, 2016) to identify specific microbial community members that represented each hydrologic treatment. Next, we examined the relationship between individual GHG fluxes and microbial community Bray-Curtis dissimilarity matrix using distance based partial least square regression in the *dbstats* R package (Boj et al., 2017). Finally, we used Mantel R statistic function in the *vegan* R package (Oksanen, 2015) to examine the relationship between microbial community composition and all soil properties (moisture, pH, organic matter, NH_4_^+^ and NO_3_^-^ concentrations, total C, total N, and C:N).

## 3. Results

### 3.1 Sediments and water properties

Over the one year study period in 2017, there were 268 days without precipitation. Within flooded plots, water depth ranged from 8-28 cm in shallow water areas and 25-53 cm in deep pools. Temperatures varied seasonally with July having the highest average (± SD) temperature (26.5 ± 1.8°C) and December having the lowest average temperature (4.9 ± 1.3°C) (Table 1). Sediments in the wetland were acidic (4.6 ± 0.3 pH) ranging from pH 4.08 to 5.28. Extractable NH_4_^+^ and NO_3_^-^ concentrations were generally low in sediments (0.089 ± 0.066 mg NH_4_^+^-N g^-1^ dry mass and 0.023 ± 0.008 mg NO_3_^-^-N g^-1^ dry mass). In the water column, NH_4_^+^ concentrations were higher (2.57 ± 19.6 mg NH_4_^+^ L^-1^) than NO_3_^-^ concentrations (0.328 ± 0.803 mg NO_3_^-^-N L^-1^) throughout the wetland, especially during winter months (9.97 ± 39.4 mg NH_4_^+^ L^-1^ and 0.769 ± 1.27 mg NO_3_^-^ L^-1^). Sediment C:N ratios along with other sediment parameters (total C, total N, and C:N) and water column phosphate (PO_4_^3-^) concentrations were similar across space (hydrologic zone) and time (season) within the wetland (Table 1A&B).

**Table 1A & B.** Summary of mean (range) of soil and water properties measured at the constructed wetland. Annual wetland values are averages of all samples for the entire sampling period. (1A) Hydrology (flooded vs. shallow land) values are averages of samples collected in May (spring), July (summer), October (fall), and December (winter) for each hydrology type. (1B) Seasonal sediment values are averages from of all plots based on samples collected in May (spring), July (summer), October (fall), and December (winter). Seasonal water values are averages of monthly samples from flooded plots where spring represents months May-June, summer represents months July-September, fall represents months October-November, and winter represents months December-February. (Abbreviations: DM = dry mass. MDL = below detection limit).

| A - Hydrology | Whole Wetland |  | Flooded |  | Shallow land |  | 345 |
| --- | --- | --- | --- | --- | --- | --- | --- |
|  | Mean | Range | Mean | Range | Mean | Range |  |
| Soil |  |  |  |  |  |  | 346 |
| temperature (°C) | 18.6 | 3.0-30.0 |  |  |  |  |  |
| moisture (%) | 33.3 | 18.3-51.9 | 36.2 | 23.2-51.9 | 31.5 | 18.3-46.2 | 347 |
| pH | 4.6 | 4.1-5.3 | 4.7 | 4.2-5.3 | 4.6 | 4.1-5.2 |  |
| organic matter (%) | 6.20 | 3.40-10.17 | 6.54 | 3.41-10.17 | 6.01 | 3.56-8.42 |  |
| mg NH<sub>4</sub><sup>+</sup>-N g<sup>-1</sup> DM | 0.089 | 0.023-0.457 | 0.129 | 0.049-0.457 | 0.066 | 0.023-0.148 |  |
| mg NO<sub>3</sub><sup>-</sup>-N g<sup>-1</sup> DM | 0.023 | 0.012-0.050 | 0.027 | 0.14-0.050 | 0.021 | 0.012-0.038 |  |
| Total C (%) | 1.66 | 0.27-3.61 | 1.78 | 0.27-3.61 | 1.60 | 0.57-2.81 |  |
| Total N (%) | 0.10 | 0.02-0.20 | 0.11 | 0.02-0.20 | 0.10 | 0.04-0.16 |  |
| C:N (wt: wt) | 15.78 | 11.00-19.00 | 15.88 | 11.00-19.00 | 15.73 | 13.88-18.44 |  |

| B - Seasonal | Spring |  | Summer |  | Fall |  | Winter |  | 348 |
| --- | --- | --- | --- | --- | --- | --- | --- | --- | --- |
|  | Mean | Range | Mean | Range | Mean | Range | Mean | Range |  |
| Soil |  |  |  |  |  |  |  |  | 349 |
| temperature (°C) | 19.6 | 16.8-23.0 | 26.5 | 25.0-30.0 | 21.3 | 20.0-24.0 | 4.9 | 3.0-7.0 |  |
| moisture (%) | 32.1 | 26.6-42.9 | 35.3 | 27.0-48.1 | 35.6 | 26.6-51.9 | 29.3 | 18.3-44.1 | 350 |
| pH | 4.7 | 4.4-5.2 | 4.6 | 4.2-5.1 | 4.5 | 4.1-5.2 | 4.7 | 4.2-5.3 |  |
| organic matter (%) | 6.32 | 5.12-8.83 | 6.26 | 4.13-10.09 | 6.68 | 3.75-10.17 | 5.39 | 3.41-8.39 | 351 |
| mg NH<sub>4</sub><sup>+</sup>-N g-1 DM | 0.085 | 0.023-0.148 | 0.089 | 0.036-0.204 | 0.098 | 0.028-0.457 | 0.082 | 0.040-0.201 |  |
| mg NO<sub>3</sub><sup>-</sup>-N g-1 DM | 0.020 | 0.016-0.027 | 0.025 | 0.017-0.038 | 0.023 | 0.014-0.034 | 0.026 | 0.012-0.050 |  |
| Total C (%) | 1.57 | 0.62-2.74 | 1.78 | 0.27-3.61 | 1.80 | 0.48-2.81 | 1.46 | 0.33-3.49 |  |
| Total N (%) | 0.10 | 0.04-0.17 | 0.11 | 0.02-0.19 | 0.11 | 0.03-0.16 | 0.09 | 0.03-0.20 |  |
| C:N (wt: wt) | 16.0 | 15.0-18.4 | 16.2 | 13.5-19.0 | 15.9 | 13.9-17.6 | 14.9 | 11.0-17.5 |  |
| Water |  |  |  |  |  |  |  |  |  |
| mg NH<sub>4</sub><sup>+</sup>-N L-1 | 0.101 | MDL-0.474 | .206 | 0.019-0.923 | 0.263 | 0.068-1.065 | 9.972 | MDL-197.441 |  |
| mg NO<sub>3</sub><sup>-</sup>-N L-1 | 0.173 | MDL-1.118 | 0.007 | MDL-0.020 | 0.169 | MDL-0.413 | 0.769 | MDL-4.055 |  |
| Total Dissolved | 0.035 | MDL-0.087 | 0.088 | 0.27-0.286 | 0.072 | 0.025-0.176 | 0.075 | MDL-0.245 |  |

### 3.2 GHG flux rates

#### 3.2.1 Methane (CH_4_)

Across the wetland, CH_4_ fluxes differed by hydrology and season (Fig. 2A). Average (± SD) CH_4_ fluxes across all seasons were lowest in shallow land plots (2.32 ± 9.33 mg CH_4_-C m^-2^ hr^-1^) and highest in flooded plots (80.80 ± 118.31 mg CH_4_-C m^-2^ hr^-1^). Within flooded plots, the highest mean CH_4_ fluxes (160.24 ± 156.28 mg CH_4_-C m^-2^ hr^-1^) were detected during summer months, and the lowest CH_4_ fluxes were detected in winter and spring months (45.02 ± 88.74 and 46.98 ± 59.40 mg CH_4_-C m^-2^ hr^-1^, respectively). Within shallow land plots, the highest mean CH_4_ fluxes (6.38 ± 16.44 mg CH_4_-C m^-2^ hr^-1^) were detected during spring months and the lowest CH_4_ fluxes were detected in winter (0.09 ± 0.42 mg CH_4_-C m^-2^ hr^-1^). The model that included hydrology and seasonal interaction explained the most variation in CH_4_ fluxes based on a _Δ_AICc score of 0 and an AICc weight of 1 (Table S1.1). In this model the interaction of hydrology and season and random effects accounted for 73% variation in CH_4_ fluxes, while the interaction alone explained 63% of that variation (Table S1.1).

**Figure 2.**
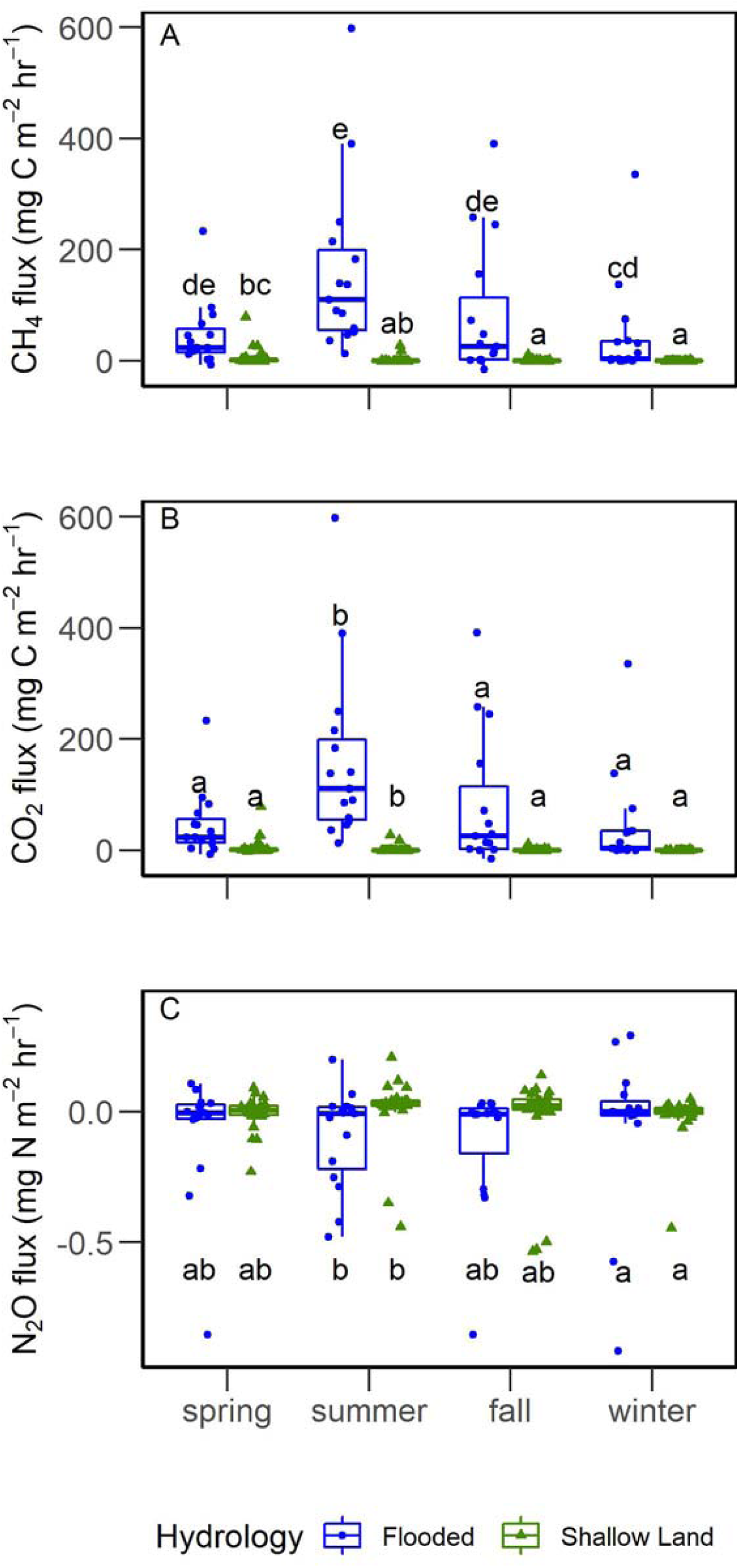
Boxplots of seasonal CH_4_ (A), CO_2_ (B), and N_2_O (C) fluxes for flooded (blue circles) and shallow land (green triangles) plots. Spring = Mar-May, Summer = Jun-Aug, Fall = Sep-Nov, Winter = Dec-Feb. Letters indicate significantly different groups based on estimated marginal means with Tukey post-hoc assessment at p<0.05 using 95% confidence levels (Table S2).

#### 3.2.2 Carbon dioxide (CO_2_)

The wetland CO_2_ fluxes varied seasonally but were generally highest in flooded plots compared to shallow land plots (Fig. 2B). Across all seasons CO_2_ fluxes were lowest in shallow land plots (3.95 ± 43.96 mg CO_2_-C m^-2^ hr^-1^) and highest in flooded plots (73.52 ± 100.24 mg CO_2_-C m^-2^ hr^-1^). We measured similar CO_2_ flux rates (20.82 ± 46.36, 14.37 ± 44.68, and 18.88 ± 24.23 mg CO_2_-C m^-2^ hr^-1^) during spring, fall, and winter seasons, while CO_2_ fluxes (63.13 ± 133.21 mg CO_2_-C m^-2^ hr^-1^) were highest during the summer. The model with season only explained the most variation in CO_2_ fluxes based on a ΔAICc score of 0 and AICc weight of 0.93 (Table S1.1). The fixed effect of season explained 26% of the variation and the full model explained 50% of the variation in CO_2_ fluxes (Table S1.1).

#### 3.2.3 Nitrous oxide (N_2_O)

The wetland N_2_O fluxes were near zero in all plots across all seasons Fig. 2C). Specifically, N_2_O fluxes were highest in shallow land plots (−0.006 ± 0.117 mg N_2_O-N m^-2^ hr^-1^) and lowest in flooded plots (−0.083 ± 0.243 mg N_2_O-N m^-2^ hr^-1^). Average N_2_O flux rates across seasons ranged from -0.05 ± 0.20 to -0.02 ± 0.15 mg N_2_O-N m^-2^ hr^-1^.

Unlike CH_4_ and CO_2_ fluxes, hydrology and season did not influence N_2_O fluxes. In this case, the null model had a ΔAICc score of 0 and the greatest AICc weight (0.69) (Table S1.1). However, the models with hydrology or season had similar ΔAICc scores (2.98, 3.98) but lower AICc weights (0.15, 0.11) when compared to the null model. Comparison of the parameters hydrology and season using estimated marginal means at 95% confidence level suggests season has the greatest impact on N_2_O fluxes (Table S1.2).

### 3.3 Global warming potential

In terms of GWP, flooded plots have a two-fold increase in magnitude of total warming potential than shallow land plots especially during warmer months (2418.63, 58.46 CO_2_ equivalents (mg m^-2^ hr^-1^), respectively) (Fig. 3). This is primarily due to increases in CH_4_ concentrations, which make up 99% of the CO_2_ equivalents in both flooded and shallow land plots (Fig. 3). Over one year, flooded areas within the wetland had the potential to produce about 7161.27 kg CO_2_ equivalents, while shallow land areas produced about 93.20 kg CO_2_ equivalents.

**Figure 3.**
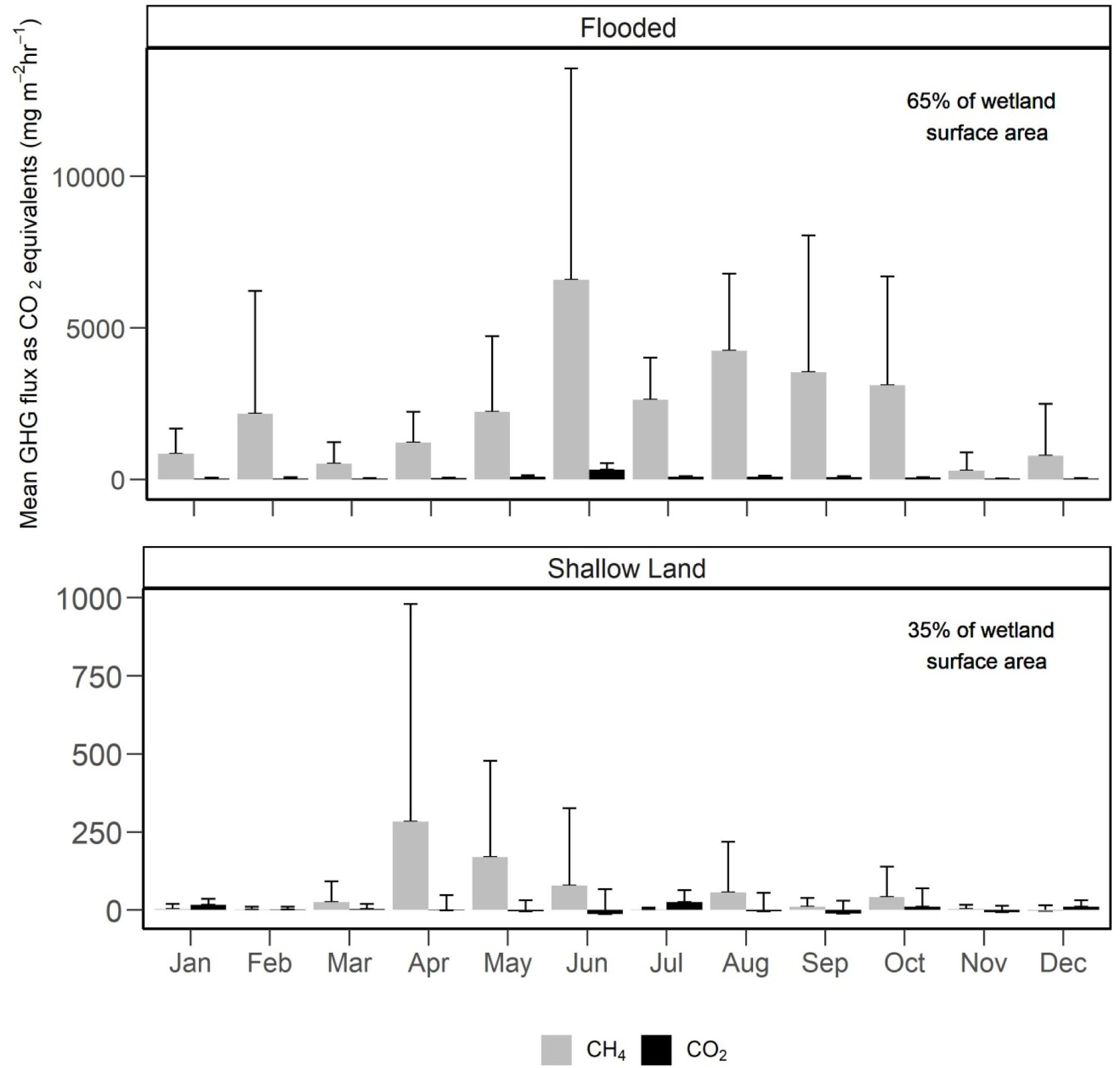
Monthly estimated global warming potentials of methane and carbon dioxide evaluated as carbon dioxide equivalents (+ standard error) measured over a year at a constructed wetland. Proportion of grey for each bar represents mg CH4-C m-2 hr-1 converted to CO2 equivalents and proportion of black for each bar represents mg CO2-C m-2 hr-1. Average mg CH4-C m-2 hr-1 was multiplied by 28, which is the estimated increased radiative force of methane when compared to CO_2_.

### 3.4 Denitrification potential

When nitrate is readily available and anoxic conditions are present, sediments from flooded and shallow land zones have similar denitrification potential (Fig. 4). In flooded and shallow land plots across all seasons, potential denitrification rates were 24.45 ± 20.18 and 20.29 ± 15.14 ng N_2_O-N hr^-1^ g^-1^ dry mass, respectively. However, denitrification potential varied by season; we measured higher rates in spring and summer (34.12 ± 21.09 and 25.16 ± 13.51 ng N_2_O N hr^-1^ g^-1^ dry mass) than in fall and winter (10.55 ± 6.17 and 16.45 ± 13.72 N_2_O ng N hr^-1^ g^-1^ dry mass) months. The model with season only explained the most variation in potential denitrification rates based on a ΔAICc score of 0 and AICc weight of 0.78 (Table S1.3). The fixed effect of season explained 18% of the variation; however, the null model explained the majority of variation (42%) (Table S1.3). We selected the season only model even though the model with hydrology and season had a similar ΔAICc (1.27) and AICc weight of 0.21, but there was little improvement in marginal and conditional R^2^ values (Table S1.3).

**Figure 4.**
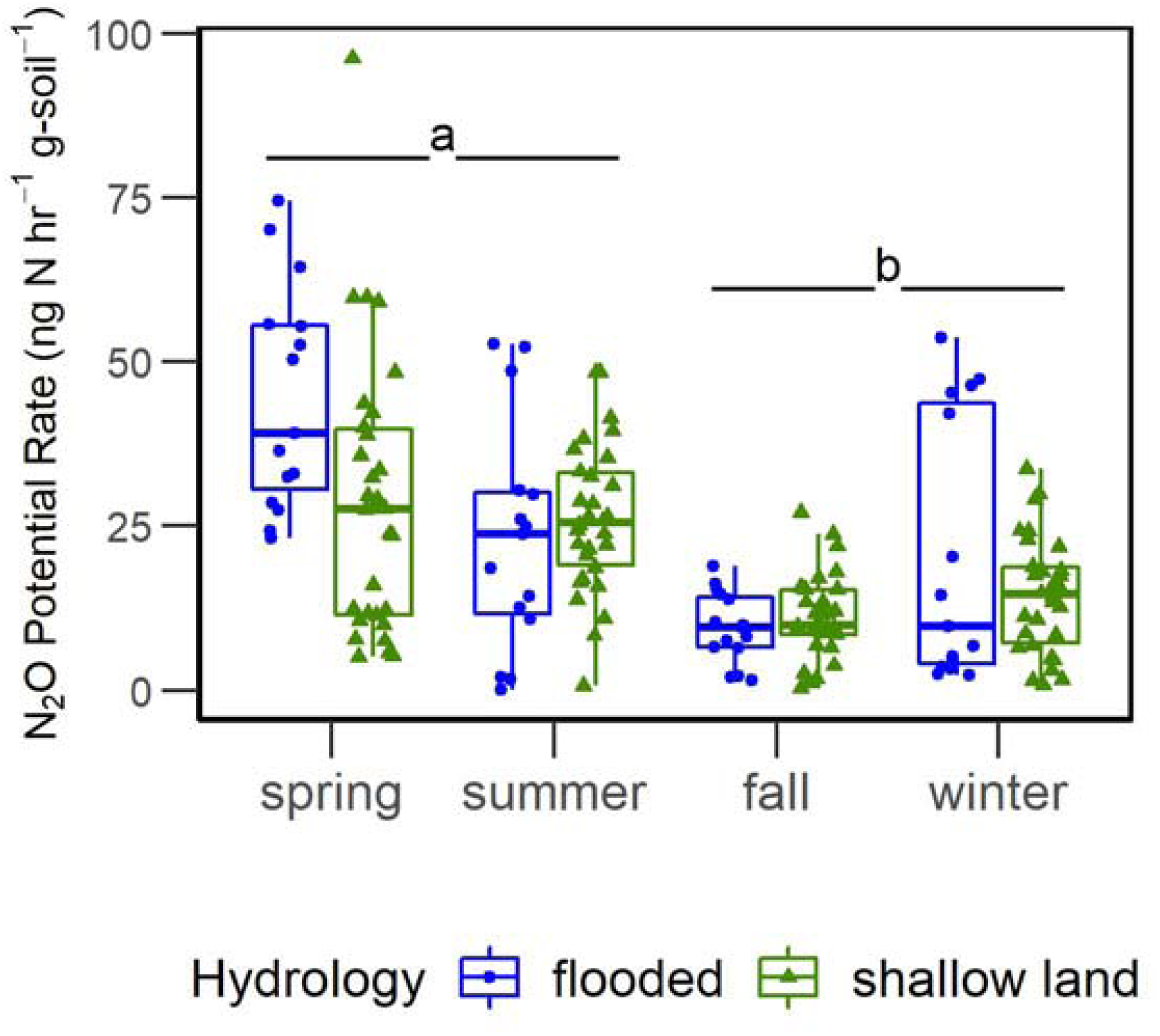
Boxplots of potential denitrification rates according to hydrology and season based on denitrification enzyme assay using the acetylene block method. Spring = Mar-May, Summer = Jun-Aug, Fall=Sep-Nov, Winter=Dec-Feb. Different letters indicate significantly different groups by Tukey adjusted estimated marginal means p<0.05. Blue circles indicate flooded plots and green triangles shallow land.

### 3.5 Microbial community analysis

Hydrology, and to a lesser extent season, influenced microbial community composition. Illumina amplicon sequencing of the 16S rRNA V4 region resulted in 454,709 reads before and 435,331 reads after removing singletons and doubletons. Each sample (n=22) then comprised a minimum of 12,510 reads and all samples were rarified to this minimum and resampled. The final dataset contained a total of 14,563 OTUs among all rarified samples. Microbial community composition in flooded plots were distinct from shallow land plots (PERMANOVA, R^2^= 0.0886, p=0.007; Fig. 5). Indicator species analysis identified one Operational Taxonomic Unit (OTU; microbial taxon defined at 97% sequence similarly), in the family Syntrophaceae, that was unique to flooded plots and identified eight OTUs that represented the shallow land plots. The shallow land indicator taxa included OTUs from two unclassified bacteria, two Betaproteobacteria, one Acidobacteria Gp6, and one *Geobacter* (Table S5). Further examination of taxa with >0.6% relative abundance of the total community suggests that flooded zones are dominated by taxa reported to be involved in methanogenic degradation of hydrocarbons (e.g., Methanomicrobia, Methylocystis, and Syntrophaceae), while shallow land zones have fewer methanogens in comparison to flooded zones (Fig. 6). We chose to look at the top 0.6% of microbial groups within the community because this threshold allowed us to further examine changes in relative abundance of the indicator species Syntrophaceae; this group is implicated in methanogenic hydrocarbon degradation and has greater relative abundance in flooded compared to shallow land areas. In addition, shallow land zones are enriched with taxa known to be involved in nitrogen cycle transformations (e.g., Bradyrhizobium, Nitrososphaera, Hyphomicrobium) (Fig. 6).

**Figure 5.**
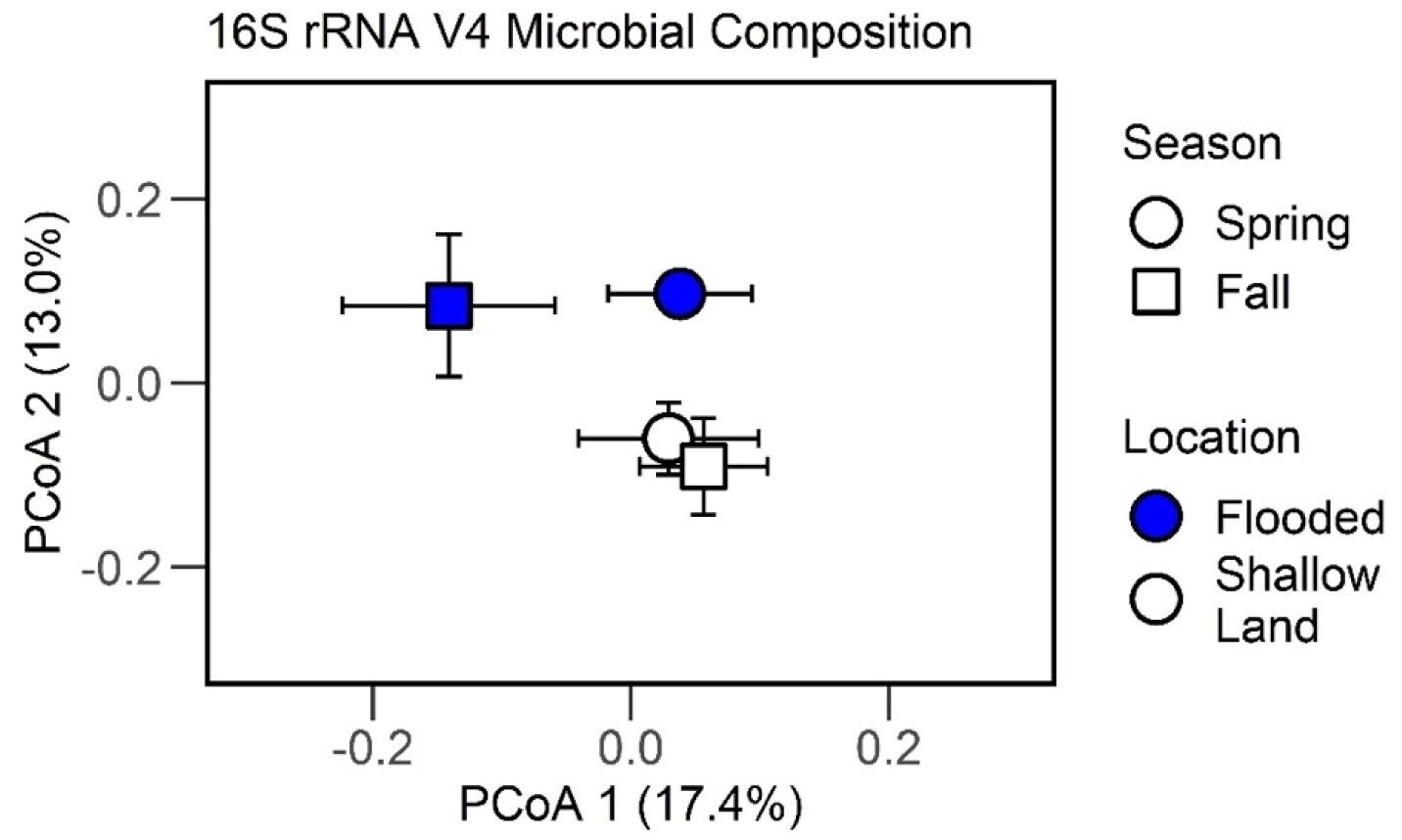
Ordination plot based on Principal Coordinates Analysis depicting sediment microbial community composition. Each point represents the centroid and range across season and sampling location. Symbols represent sampling period (circle = spring, square = fall). Spring samples were collected in May and fall samples were collected in October. Colors represent sampling plots (blue = flooded, white = shallow land).

**Figure 6.**
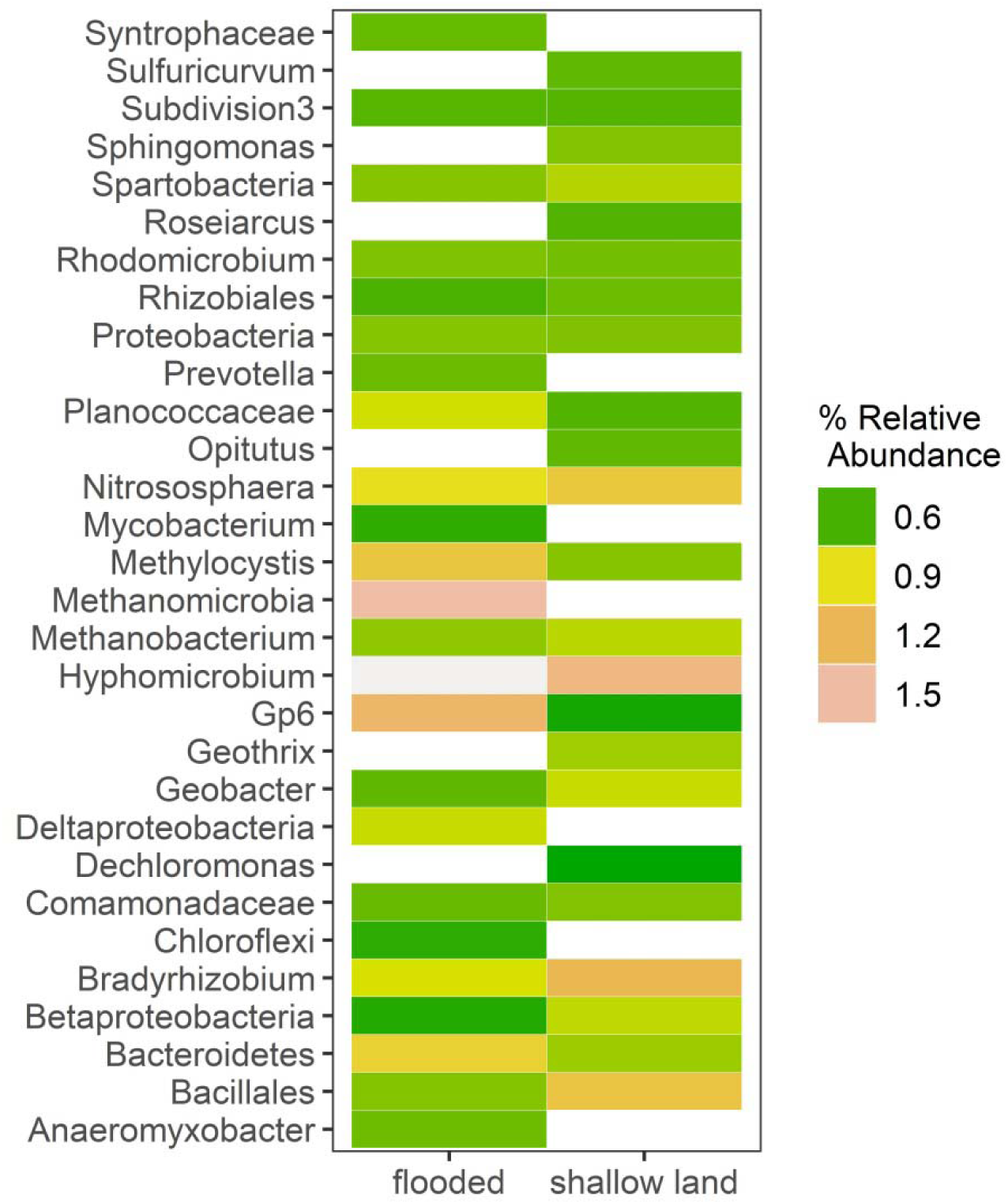
Heat map of bacterial and archaeal taxa observed at >0.6% relative abundance. Color gradient (cool to warm, green to red) represents microbial relative abundance ∼0.6 to 1.5%.

### 3.6 Analyses of relationship between GHGs, sediment chemistry, and microbial community composition

Sediment chemical properties shape microbial community composition and microbial community composition influences GHG fluxes more than sediment chemistry properties. Distance-based partial least square regression analysis suggests that microbial community composition explained 82.4%, 78.6%, and 79.1%, of variation in CH_4_, CO_2_, and N_2_O fluxes, respectively, by components 1 and 2. Sediment chemistry components 1 and 2 best explained variation in N_2_O production (51.4%) but less so for CH_4_ or CO_2_ production (21.0%, 16.6%). Mantel correlation analyses revealed a positive relationship between patterns in microbial community composition and soil properties (r = 0.52, p = 0.003).

## 4. Discussion

Examination of microbial processes occurring at permanently and periodically inundated locations within a stormwater CW revealed that modifications to wetland design could enhance water quality (via denitrification) and reduce GHG production. Other studies comparing different types of CWs indicate that free water surface (FWS) wetlands, like this study site, are known to produce copious amounts of GHGs compared to vertical or horizontal subsurface flow CWs (Mander et al., 2014; McPhillips and Walter, 2015). While vertical or subsurface CWs may reduce GHG production, this type of installation is more costly than FWS wetlands and may not always be practical. We recognize that this study focuses on a single CW; however, our results demonstrate the need for more comprehensive studies exploring CW design features that can be modified to enhance beneficial microbial ecosystem services (i.e., nitrogen removal) while reducing ecosystem disservices (i.e., GHG emissions).

At our focal CW, CH_4_ was the dominant GHG produced in flooded plots across all seasons. Hydrologic conditions accounted for the most variation in CH_4_ fluxes and microbial community composition. Since high NO_3_^-^ concentrations suppress methanogenesis (Kim et al., 2015), it is likely the combination of anoxic conditions (McPhillips and Walter, 2015), low NO_3_^-^ concentrations in sediments and water, and availability of organic carbon in flooded plots provided optimal conditions for CH_4_ production (Rahman et al., 2019). However, season and microbial community composition were stronger predicators than hydrology and distance from inlet of CO_2_ and N_2_O rates within the wetland. We propose that season and hydrology played a stronger role than distance from inlet in GHG fluxes and denitrification potential due to the small size of the study CW. Seasonal differences in temperature and vegetation status (i.e., organic carbon inputs) within this study influenced CO_2_ and N_2_O rates to a greater degree than hydrology. In other studies, temperature and substrate availability strongly determined rates of organic decomposition and denitrification (Davidson and Janssens, 2006; Knowles, 1982; Moinet et al., 2018).

Overall, compared to wastewater treatment wetlands, flooded plots within our study wetland represent a considerable source of GWP primarily due to CH_4_ emissions. In order to simultaneously compare the biogenic GHGs within the CW, we quantified GWP associated with flooded and shallow land zones. In terms of GWP, CH_4_ from flooded plots was the primary carbon source within the wetland. While both CO_2_ and CH_4_ production was greatest in flooded zones compared to shallow land zones, CH_4_ production greatly exceeded CO_2_ production leading to high GWP within the wetland. Compared to surface flow wastewater treatment wetlands receiving municipal or agricultural runoff, CO_2_ emissions at our study wetland in flooded plots were similar but approximately 95% lower in shallow land plots (Jahangir et al., 2016; Mander et al., 2014). In contrast, CH_4_ emissions at our study wetland had similar CH_4_ rates in shallow land plots when compared to wastewater treatment wetlands, but approximately 94% greater emissions in flooded plots compared to wastewater treatment wetlands (Jahangir et al., 2016; Mander et al., 2014). In terms of GWP, flooded plots in our study wetland have a GWP that is 98% greater than in wastewater treatment wetlands. The increased CH_4_ emission in our study wetland compared to wastewater wetlands is likely due to the lower availability of nutrients in surface water and sediments.

Further, GHG emissions could be partially explained by microbial community structure. This is due to hydrologic conditions in flooded zones providing optimal habitat for obligate anaerobes that participate in methanogenesis. Methanogenesis is suppressed by even low oxygen concentrations (Fetzer et al., 1993; Liu et al., 2008) and we could not detect methanogens in shallow land areas where conditions can fluctuate between oxic and anoxic environments. Additionally, analysis of microbial community structure suggests that hydrocarbon degradation, a potential microbial ecosystem benefit, may be occurring in flooded plots. Hydrocarbons are becoming recognized as a common contaminant in urban wetlands with the potential to reduce and even harm wildlife taking refuge in these habitats (Clevenot et al., 2018; Mahler et al., 2014). This finding demonstrates the value of examining the microbial community composition when evaluating beneficial wetland ecosystem functions.

In retrospect of this study, we highlight three CW features for consideration to reduce ecosystem function disservices: (i) surface area of hydrologic zones, (ii) incoming nutrient and pollutant loads, and (iii) the anticipated number of rain days. Results from this study and others indicate that areas that fluctuate between oxic and anoxic conditions, which occur in CW shallow land areas in this study, are sites of reduced CH_4_ production. This is due to the inhibition of methanogenesis and a greater abundance of methanotrophs which consume methane (Chowdhury and Dick, 2013; Lew and Glińska-Lewczuk, 2018). Additionally, even in CWs with low nitrate availability, shallow land areas could potentially support enhanced denitrification due to coupling of nitrification, which occurs during oxic periods and with steady organic carbon inputs from vegetation (Mander et al., 2014; Rahman et al., 2019). Flooded zones increase water storage capacity and provide habitat for predator species that control nuisance species; therefore, elimination of flooded zones may not be desirable. However, an increase in shallow land area and a reduction in surface area of flooded zones could reduce GHG emissions without sacrificing benefits of flooded zones.

Secondly, substrate availability can directly impact microbial process rates. Compared to the neighboring Tar River (0.52 mg NO_3_-N L^-1^), seasonal NO_3_^-^ concentrations within the surface water of the CW were much lower (0.007-0.173 mg NO_3_-N L^-1^) except during winter (0.769 mg NO_3_-N L^-1^) (Humphrey et al., 2019). This increase in nitrate during winter is likely due to the senescence of *Typha* spp. and *P. cordata* vegetation (Bachand and Horne, 2000), which are the main plants growing in and around the main channel. Considering that NO_3_^-^ availability limits denitrification, ecosystems that receive runoff from agriculture, livestock, or are adjacent to a high density of septic systems are areas of high denitrification potential and low methane production (Naylor et al., 2018). Recent studies show that iron (II) (Fe^2+^) additions can increase N removal from vertical and horizontal subsurface CWs (Song et al., 2016; Zhang et al., 2019). In the case of existing wetlands, further investigation into amendments as a way to increase denitrification potential is warranted. Additionally, depending on the upstream runoff source, pollutants such as hydrocarbons may be better targets for remediation within intentionally designed CWs.

Lastly, we suggest that the frequency of rain days or inter-event duration is taken into account during SCM planning (Andersen et al., 2017). Our results represent ambient conditions, at least 48 hours after a storm event, which represent more than half the year for this CW. Within this particular wetland, reducing the permanently flooded surface area could reduce GHGs while still providing N removal benefits. In areas where rain events are less frequent (i.e., greater inter-event durations) and more intense, SCMs that are not permanently flooded, such as dry infiltration basins or dry detention basins, could provide relief for intense runoff events with reduced GHG production (McPhillips and Walter, 2015; Morse et al., 2017). Therefore, we suggest frequency of rain events as another factor to consider when planning SCM design.

## 5. Conclusion

Context matters when designing CWs in urban watersheds. This study demonstrates the importance of considering microbial controls on biogeochemical processes within SCMs in order to reduce tradeoffs between water quality and GHG production. In the case of this urban stormwater CW, which is located in a parking lot of a highly urbanized area, we suggest that reducing flooded areas and increasing shallow land area would reduce GHG emissions while still providing nutrient removal benefit. However, our analysis of the microbial community also indicates that potential hydrocarbon decomposition could be supported in flooded zones and warrants a deeper investigation of this potential service. This study also highlights the need for a comprehensive study of CW design features to better understand which design features and when those features provide the most ecosystem benefits. By examining both microbial process rates and the microbial community composition, we can approach SCM design and implementation in a holistic way that accounts for ecosystem services and disservices.

## Abbreviations

AICc: corrected Akaike information criteria,
NH_4_^+^: ammonium,
C: carbon,
CO_2_: carbon dioxide,
CW: constructed wetland,
N_2_: dinitrogen,
GWP: global warming potential,
GHG: greenhouse gas,
CH_4_: methane,
N: nitrogen,
NO_3_^-^: nitrate,
NO: nitric oxide,
NO_2_^-^: nitrite,
N_2_ O: nitrous oxide,
OTU: operational taxonomic unit,
PCR: polymerase chain reaction,
PERMANOVA: permutated analysis of variance,
PCoA: principal coordinates analysis,
SCM: stormwater control measure

## Declaration of Competing Interest

None.

## Acknowledgements

We thank B. Hinckley, A. Garcìa, and L. Armstrong for assistance with field research and sample processing. We also thank Sound Rivers for installation of the constructed wetland and ECU Grounds personnel and manager J. Gill for site maintenance. We also thank E. Field, L. Kinsman-Costello, M. McCoy, and anonymous reviewers for feedback on earlier versions of this manuscript. This work was supported by East Carolina University and the North Carolina Sea Grant/Water Resources Research Institute Graduate Student Research Fellowship and the National Science Foundation Graduate Research Fellowship to RBB. All code and data used uyhin this study can be found in a public GitHub repository (DOI: 10.5281/zenodo.3924697) and the NCBI SRA (BioProject PRJNA613095).

## Appendix A. Supplementary data

Supplementary material related to this article can be found at: ###

## Notes

https://github.com/PeraltaLab/ConstructedStormwaterWetland

### Competing Interest Statement

The authors have declared no competing interest.

### Summary of Updates

Clarifications throughout text based on reviewers suggestions. Error bars added to figure 3.

