## Supplemental Data for "A microbial perspective on balancing trade-offs in ecosystem functions in a constructed stormwater wetland" (from another version of this preprint: version 1, 2020-04-03)

*Corresponding author

**Affiliations:**

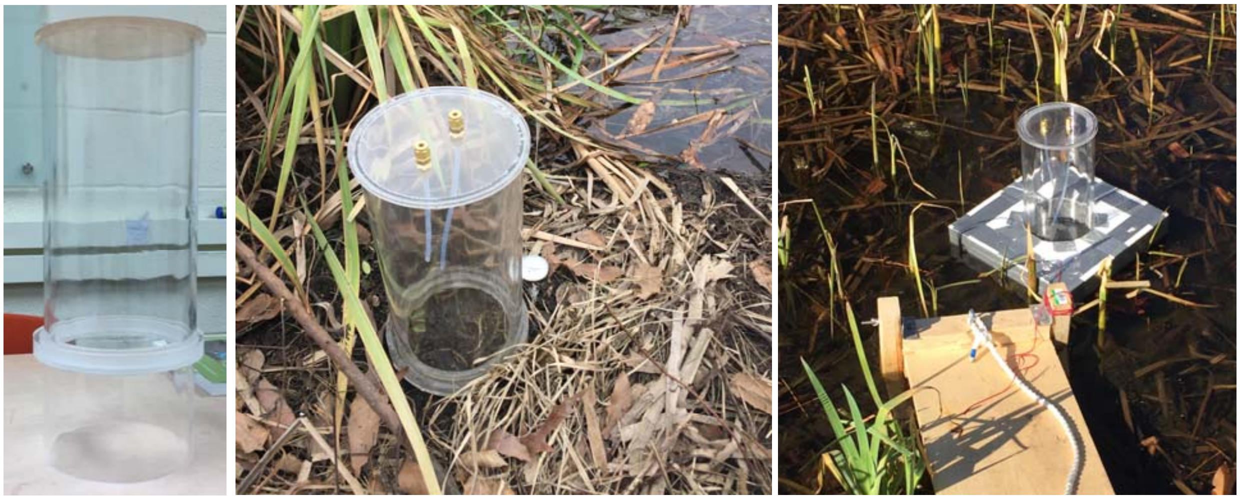

.

Figure S1. Images of GHG chambers. Left: Chamber top and bottom. Middle: Shallow land chamber installed on sampling plot. Right: Flooded plot chamber on Styrofoam float. Access boards were set up prior to sampling to allow access into flooded plots.

Table S1. Mixed-effects models for GHG fluxes with sample plot as a random effect. H = hydrology, S = season, D = distance from inlet.

| Model | k | AICc | ΔAICc | AICc Wt | $R^2$ Marg | $R^2$ Cond |
|---|---|---|---|---|---|---|
| CH4 | | | | | | |
| H*S | 10 | 313.70 | 0 | 1 | 0.63 | 0.73 |
| Hydrology | 4 | 338.78 | 25.07 | 0 | 0.55 | 0.65 |
| H*D | 9 | 342.44 | 28.73 | 0 | 0.60 | 0.64 |
| Season | 6 | 343.41 | 29.70 | 0 | 0.05 | 0.69 |
| H*D*S | 30 | 344.69 | 30.69 | 0 | 0.70 | 0.75 |
| Null | 3 | 361.39 | 47.68 | 0 | - | 0.64 |
| Distance | 6 | 366.64 | 52.94 | 0 | 0.05 | 0.64 |
| CO2 | | | | | | |
| Season | 6 | 45.54 | 0 | 0.93 | 0.26 | 0.50 |
| H*S | 10 | 50.71 | 5.16 | 0.07 | 0.46 | 0.50 |
| Hydrology | 4 | 68.40 | 22.86 | 0 | 0.22 | 0.23 |
| Null | 3 | 78.06 | 32.51 | 0 | 0 | 0.24 |
| H*D | 9 | 86.67 | 41.12 | 0 | 0.26 | 0.26 |
| Distance | 6 | 87.37 | 41.82 | 0 | 0.07 | 0.28 |
| H*D*S | 29 | 109.20 | 63.65 | 0 | 0.56 | 0.56 |
| N2O | | | | | | |
| Null | 3 | 264.19 | 0 | 0.69 | 0 | 0.17 |
| Hydrology | 4 | 267.17 | 2.98 | 0.15 | 0.02 | 0.19 |
| Season | 6 | 267.88 | 3.69 | 0.11 | 0.05 | 0.23 |
| Distance | 6 | 270.11 | 5.93 | 0.04 | 0.07 | 0.20 |
| H*D | 9 | 271.91 | 7.72 | 0.01 | 0.08 | 0.26 |
| H*S | 10 | 278.96 | 14.77 | 0 | 0.17 | 0.21 |
| H*D*S | 30 | 316.91 | 52.72 | 0 | 0.27 | 0.32 |

Table S2. GHG flux estimated marginal means at 95% confidence levels (CL) with Tukey (alpha=0.05) adjusted post-hoc analysis of the selected model parameter based on AICc analysis. Log cuberoot transformed values. SL = shallow land, F = flooded, W = winter, FA = fall, SM = summer, SP = spring

| Parameter | emmean | SE | df | lower.CL | upper.CL | group |
|---|---|---|---|---|---|---|
| CH4: interaction | | | | | | |
| SL*W | -0.828 | 0.162 | 44.3 | -1.154 | 0.501 | a |
| SL*FA | -0.725 | 0.159 | 41.4 | -1.046 | -0.404 | a |
| SL*SM | -0.634 | 0.159 | 41.2 | -0.955 | -0.314 | ab |
| SL*SP | -0.191 | 0.160 | 42.8 | -0.515 | 0.132 | bc |
| F*W | 0.574 | 0.221 | 39 | 0.127 | 1.021 | cd |
| F*FA | 0.949 | 0.221 | 39.0 | 0.503 | 1.396 | de |
| F*SP | 1.085 | 0.221 | 39.0 | 0.638 | 1.532 | de |
| F*SM | 1.554 | 0.221 | 39.0 | 1.107 | 2.001 | e |
| CO2: Season | | | | | | |
| Winter | 0.602 | 0.0468 | 57.1 | 0.508 | 0.696 | a |
| Spring | 0.743 | 0.0473 | 58.4 | 0.648 | 0.837 | a |
| Fall | 0.744 | 0.0505 | 63.8 | 0.643 | 0.845 | a |
| Summer | 1.059 | 0.0481 | 58.3 | 0.963 | 1.156 | b |
| N2O: Hydrology | | | | | | |
| F | -1.29 | 0.089 | 13.4 | -1.48 | -1.094 | a |
| SL | -1.13 | 0.123 | 12.2 | -1.40 | -0.862 | a |
| Season | | | | | | |
| Winter | -1.42 | 0.101 | 46.1 | -1.62 | -1.218 | a |
| Spring | -1.28 | 0.100 | 46.1 | -1.48 | -1.078 | ab |
| Fall | -1.18 | 0.098 | 42.6 | -1.38 | -0.986 | ab |
| Summer | -1.06 | 0.099 | 43.7 | -1.26 | -0.861 | b |

Table S3. Mixed-effects models for DEA $N_2O$ flux potentials with sample plot as a random effect. H = hydrology, S = season, D = distance from inlet.

| Model | k | AICc | ΔAICc | AICc Wt | $R^2$ Marg | $R^2$ Cond |
|---|---|---|---|---|---|---|
| H*S | 10 | 478.74 | 0 | 0.65 | 0.21 | .047 |
| Season | 6 | 480.01 | 1.27 | 0.35 | 0.18 | 0.42 |
| H*D*S | 18 | 513.69 | 34.95 | 0 | 0.38 | 0.55 |
| Null | 3 | 516.17 | 37.42 | 0 | - | 0.23 |
| Distance | 6 | 518.36 | 39.62 | 0 | 0.08 | 0.27 |
| Hydrology | 4 | 518.68 | 39.94 | 0 | 0.00 | 0.25 |
| H*D | 6 | 529.25 | 50.50 | 0 | 0.10 | 0.26 |

Table S4. DEA $N_2O$ flux potentials estimated marginal means at 95% confidence levels (CL) with Tukey (alpha=0.05) adjusted post-hoc analysis of the selected model parameter based on AICc analysis. Log cuberoot transformed values. SL = shallow land, F = flooded, W = winter, FA = fall, SM = summer, SP = spring

| Parameter | emmean | SE | df | lower.CL | upper.CL | group |
|---|---|---|---|---|---|---|
| Interaction | | | | | | |
| F*FA | 2.03 | 0.322 | 26.7 | 1.37 | 2.69 | ab |
| SL*FA | 2.16 | 0.229 | 27.4 | 1.69 | 2.63 | a |
| SL*W | 2.37 | 0.227 | 26.7 | 1.91 | 2.84 | a |
| F*W | 2.43 | 0.322 | 26.7 | 1.77 | 3.09 | ab |
| F*SM | 2.52 | 0.322 | 26.7 | 1.86 | 3.18 | ab |
| SL*SP | 3.08 | 0.227 | 26.7 | 2.61 | 3.55 | bc |
| SL*SM | 3.10 | 0.227 | 26.7 | 2.63 | 3.57 | bc |
| PI*SP | 3.73 | 0.322 | 26.7 | 3.07 | 4.39 | c |
| Season | | | | | | |
| Fall | 2.12 | 0.184 | 31.7 | 1.74 | 2.49 | a |
| Winter | 2.39 | 0.183 | 31.1 | 2.02 | 2.76 | a |
| Summer | 2.91 | 0.183 | 31.1 | 2.53 | 3.28 | b |
| Spring | 3.29 | 0.183 | 31.1 | 2.92 | 3.67 | b |

Table S5. Summary of bacterial taxa (OTUs) characteristic to flooded and shallow land plots based on indicator species analysis.

These are the top OTUs that are significantly associated with each hydrology.

| OTU | Cluster | IndVal | Prob | Phlyum | Order | Class | Family | Genus |
|---|---|---|---|---|---|---|---|---|
| Otu00037 | F | 0.730 | 0.002 | Proteobacteria | Deltaproteobacteria | Syntrophobacterales | Syntrophaceae | unclassified |
| Otu00007 | SL | 0.632 | 0.002 | Verrucomicrobia | Spartobacteria | unclassified | unclassified | unclassified |
| Otu00010 | SL | 0.604 | 0.006 | Bacteria | unclassified | unclassified | unclassified | unclassified |
| Otu00013 | SL | 0.630 | 0.007 | Acidobacteria | Acidobacteria_Gp6 | Gp6 | unclassified | unclassified |
| Otu00014 | SL | 0.576 | 0.02 | Proteobacteria | Betaproteobacteria | unclassified | unclassified | unclassified |
| Otu00025 | SL | 0.680 | 0.006 | Proteobacteria | Betaproteobacteria | unclassified | unclassified | unclassified |
| Otu00031 | SL | 0.597 | 0.033 | Bacteria | unclassified | unclassified | unclassified | unclassified |
| Otu00032 | SL | 0.596 | 0.019 | Proteobacteria | Deltaproteobacteria | Desulfuromonadales | Geobacteraceae | Geobacter |
